# Sleep dynamics and epileptogenesis following Kainic acid in epilepsy-susceptible (DBA/2J) and epilepsy-resistant (C57BL/6) mice

**DOI:** 10.64898/2026.01.16.699940

**Authors:** Jun Wang, Jesse Isaacson, Anna Goforth, Rama Maganti

## Abstract

Susceptibility to epileptogenesis varies in humans and outbred mouse strains. We hypothesized that baseline sleep abnormalities increase susceptibility to epileptogenesis following Kainic acid (KA) and abnormal circadian rhythm or sleep homeostasis (SH) contribute to worse seizures in epilepsy-susceptible DBAs. Following EEG electrode implantation, C57 and DBA mice underwent repeated low-dose KA or saline treatment. Seizures, interictal spikes and sleep were examined over 8-weeks with continuous electroencephalography (EEG). Seizures were manually scored, and interictal spikes and sleep were analyzed using machine learning algorithms. Slow wave activity (SWA) was derived from non-rapid eye movement (NREM) sleep following Fourier transform, and SH was measured by the decay of SWA during sleep and its rise with preceding wakefulness. The impact of post-KA seizures on circadian rhythm was determined using Cosinor analysis. Seizures were longer and more frequent in KA-treated DBAs than C57s. Interictal spike were much greater in saline-treated DBAs than C57s. SWA across the 24-hours was lower in DBAs at baseline. KA treatment decreased REM and increased SWA activity in DBAs but not in C57s. Cosinor analysis revealed circadian rhythm abnormalities in DBAs but not in C57s. Seizures impaired SH in DBAs, with no increase in SWA with preceding wakefulness and a progressive loss of SWA decay during lights-on over the 8-week recording. These findings suggest that baseline sleep abnormalities, poorly adaptable circadian rhythms and impaired SH are associated with increased vulnerability to epileptogenesis. Therapies enhancing circadian rhythm and SH after an insult may be avenues to mitigate epileptogenesis in vulnerable populations.

**Significance Statement:** Susceptibility to spontaneous seizures after a central nervous system insult varies in humans as well as mouse models. The reasons behind the differential susceptibility are not entirely known. Here we show in a Kainic acid model, epilepsy susceptible DBA/2J mice have baseline sleep abnormalities and following Kainic acid treatment, these mice develop more frequent and longer seizures than the epilepsy-resistant C57BL/6 mice. We also show that the epilepsy-susceptible DBAs have abnormalities in circadian rhythm as well as sleep homeostasis measures compared to epilepsy-resistant C57s, which may be contributing to progression of epileptogenesis and worse seizures. Therapies targeted at enhancing sleep, circadian rhythms or sleep homeostasis may be avenues to mitigate development of spontaneous seizures after a central nervous system insult.

## Introduction

Epilepsy is a disorder characterized by recurrent seizures that affects >50 million people worldwide (Feigin et al, 2021). To understand the pathophysiology and basic mechanisms of epilepsy, a variety of rodent models have been developed over the last few decades. Among these, kindling models and chemoconvulsant models are the most commonly used (Löscher, 2011). Within chemoconvulsant models, Kainic acid (KA) is employed as the spectrum of pathophysiology and seizure phenotypes it induces in rodents closely reflects the pathophysiology of the most common human epilepsy, namely temporal lobe epilepsy (Rusina et al, 2021). However, susceptibility to develop spontaneous seizures following KA varies across mouse strains. Among all inbred mouse strains, C57BL/6 mice exhibit low seizure sensitivity compared to other strains though this is based on electroconvulsive threshold data. Because of this, C57s are termed epilepsy-resistant whereas other strains such as FVB, DBA and C3H are considered to be epilepsy-susceptible (Frankel et al, 2001, Löscher et al, 2017). Moreover seizure-induced cell death and seizure induced mortality varies among mouse strains (Shauwecker 2012). Susceptibility to epilepsy is similarly variable in humans following a central nervous system injury (Pease et al, 2022). The underlying reasons for these differences in seizure susceptibility—whether in mouse strains or in humans—are not fully understood, although genetic influences were suggested (McKhann et al, 2003).

Sleep disturbances are common in both humans and mouse models of epilepsy (Matos et al, 2010). Reported sleep disturbances in these contexts include reduced REM sleep and decreased sleep efficiency (Calvello et al, 2023). Increased sleep slow wave activity (SWA) has been observed in humans with focal epilepsy (Boly et al, 2017) and in some mouse models of epilepsy (Konduru et al, 2021; Wang et al, 2025). However, whether sleep disruptions are antecedent to the development of chronic spontaneous seizures remains unknown. Obstructive sleep apnea, a common sleep disorder that causes sleep disruption, has been linked to epilepsy risk. In older adults, the presence of obstructive sleep apnea and hypoxia has been associated with late-onset epilepsy, suggesting that sleep disturbances of any origin may contribute to de novo epilepsy (Carosella et al, 2024). Currently, there are no data available to elucidate how sleep changes evolve during epileptogenesis. In this study, we examined sleep characteristics, including sleep homeostasis, in epilepsy-resistant (C57BL/6) and epilepsy-susceptible (DBA/2J) mouse strains both prior to and following an epileptogenic insult induced by kainic acid (KA).

Sleep homeostasis (SH) is a regulatory process that maintains appropriate sleep duration and intensity by exerting a hypothetical “sleep pressure,” which builds during wakefulness to promote sleep and dissipates during sleep to permit waking (Boberly and Acherman, 1999). Experimentally, sleep pressure is quantified by measuring power in the delta frequency band (0.5–4 Hz), or slow wave activity (SWA), in EEG recordings during non-rapid eye movement (NREM) sleep. In normal subjects, delta power increases upon entry into NREM sleep following prolonged wakefulness, reflecting the buildup of sleep pressure. Conversely, delta power gradually declines with sustained sleep, indicating the dissipation of the sleep pressure (Franken and Dijk, 2024). There is limited data in the literature on evolution of spontaneous recurrent seizures and sleep changes following KA using chronic continuous Video-EEG recordings. Here, we demonstrate that DBA/2J mice, exhibit reduced baseline sleep SWA, indicative of diminished sleep homeostatic drive. Using 8-week long EEG recordings, we show that DBA mice express more frequent and longer seizures, have abnormal circadian rhythms and impaired sleep homeostasis that progressively worsen through epileptogenesis compared to C57s. We hypothesize that mouse strain with sleep abnormalities have increased susceptibility to seizures post-KA and have worse seizures that further disrupt sleep homeostasis and circadian rhythms.

## Methods

### Animals

Twelve-week-old C57BL/6J and DBA/2J mice were obtained from Envigo (Indianapolis, IN) and Jackson Laboratories (Bar Harbor, ME), respectively. All procedures involving the use of animals were approved by the University of Wisconsin Institutional Animal Care and Use Committee (IACUC) and conducted in accordance with the U.S. Department of Agriculture Animal Welfare Act and the National Institutes of Health guidelines for the Humane Care and Use of Laboratory Animals.

### EEG implantation, KA treatment and EEG recordings

Under sterile conditions and isoflurane anesthesia, all mice underwent surgical implantation of a right frontal and a left parietal EEG electrode, along with an occipital reference electrode and braided EMG leads in the nuchal muscles. Mice were allowed to recover for two days postoperatively before EEG recordings commenced. EEG and EMG signals were recorded using an XLTEK EEG system (Natus, Middleton, WI) at a 512 Hz sampling rate and bandpass filtered between 1 and 35 Hz for EEG and 10-250 Hz for EMG. Recordings were conducted 24 hours a day for 5 consecutive days each week over a period of 8 weeks. During recording sessions, mice were singly housed in 10 × 6-inch cylindrical chambers, connected to the EEG system via a tether attached to an implanted headcap. When not recording, mice were group housed. All animals were maintained on a 12:12-hour light:dark cycle (lights-on at 6:30 AM and off at 6:30 PM), with ad libitum access to food and water.

A baseline day of EEG recording was performed prior to KA administration in a subset of mice (n = 5 per group for C57 and DBA). Subsequently, mice were assigned to receive repeated low-dose saline (C57 n=15; 7 male and 5 female- and DBAs n=10; 6 male and 4 female) or KA injections (C57 n=25; 15 female and 10 female; and DBA n=18; 10 female and 8 male), via intraperitoneal (IP) administration. KA-treatment protocol consisted of initial 10 mg/kg bolus followed by alternating doses of 5 mg/kg and 2.5 mg/kg every 20 minutes until onset of status epilepticus (SE), defined as two Racine class 5 seizures within a 20-minute period. The latency to SE varied between individual animals, requiring between 5 and 15 KA injections to reach this criterion. Control mice received equivalent volumes of SA every 20 minutes. Mortality was observed in 10% of C57s and 30% DBAs. Following treatment, mice were allowed to recover spontaneously, provided with gel food and water ad lib. Continuous EEG recordings were then performed.

### Seizure analysis and interictal spike detection

Seizures were manually detected and interictal spikes (Figure 1A-C) were detected using a custom MATLAB script from the frontal EEG. Each EEG recording (in EDF format) was first bandpass filtered between 0.5 and 30 Hz. Zero-crossing points were then identified to segment the EEG signal into consecutive waveforms. A spike was defined as a complete biphasic waveform, consisting of three consecutive zero crossings: a positive wave followed by a negative wave. Events were classified as interictal spikes if they met all of the following criteria: (1) peak-to-peak duration shorter than 150 ms, (2) negative wave duration longer than 50 ms, (3) negative peak amplitude less than six times the standard deviation of the full filtered EEG signal, (4) total peak-to-peak amplitude greater than eight times the standard deviation, and (5) separated by at least 2 seconds between them (to avoid inclusion of any spike wave discharges which are common in DBAs especially-Letts et al, 2014)

**Figure 1:**
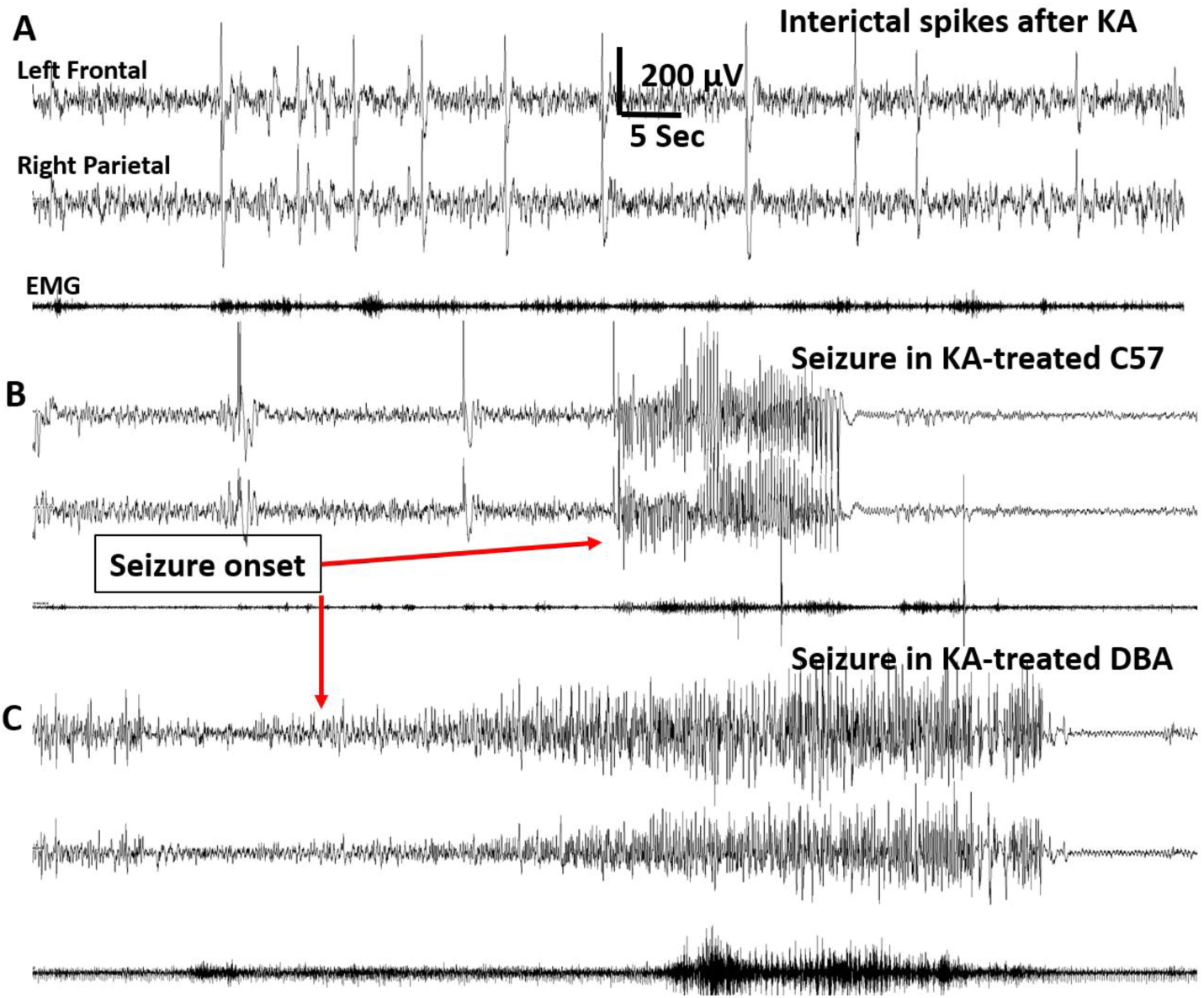
Examples of interictal spikes and seizures: A: Interictal spikes example shown from a DBA mouse. B: Seizure recorded in a C57 and C: Seizure recorded in a DBA mouse. Note that seizures are much shorter in duration in the C57 mouse.

### Sleep classification

European data format (EDF) of the EEG was used for sleep stage classification using an automated classifier made in-house called REST (Wang et al, 2025). Briefly, the short-time Fourier transforms (STFT) of frontal EEG and EMG signals were used as input features. These spectrograms were processed through two stacked Transformer encoder layers to extract both the frequency characteristics within each epoch and the temporal dependencies across consecutive epochs. The Transformer’s self-attention mechanism enabled the model to weigh the relevance of surrounding epochs, improving continuity and accuracy in stage transitions. Finally, the model assigned a sleep stage label to each epoch based on the weights learned during training, producing classifications that closely align with manual scoring.

### Sleep and SWA analysis

After excluding EEG recordings with poor signal quality (overwhelming artifact, noise, loss of signal) those suitable for analysis were then separated by mouse strain, treatment group, and seizure occurrence into 6 groups (baseline C57 and DBA n=5 days each; C57 saline n=11, C57 KA without seizure n=6, C57 KA with seizure n=9, DBA saline n=6, DBA without seizure n=8, and DBA with seizure n-6; n-recorded days=440,184,360,216,278,172). Sleep metrics including SWA were then calculated from recordings that were scored by REST. SWA is defined as the power in the delta band during NREM sleep that was normalized by dividing with the sum power in other bands (Theta, Alpha, Beta, and Gamma). The sleep metrics calculated include percentage time and mean bout duration of wake, NREM and REM sleep.

### Power spectral Density analysis

To examine frequency-specific differences in NREM sleep epochs between experimental groups, power spectral density (PSD) was computed for each animal. Raw EEG signal was first notch-filtered at 60 Hz to remove powerline interference. The PSD of each epoch was estimated using Welch’s method (scipy.signal.welch) with 0.25 Hz frequency resolution without overlapping. For each recording, the resulting spectra were truncated to 0–20 Hz. Within each animal, all NREM epoch spectra from multiple weeks of recording were averaged to yield a single mean spectrum. To allow comparison across animals, each mean spectrum was then z-scored along the frequency dimension. These normalized spectra were then grouped by experimental conditions.

### Cosinor analysis

To examine the circadian modulation of SWA and sleep stage transitions before and after the first seizure, we performed a cosinor-based mixed-effects analysis. EEG data were segmented into twelve 2-hour bins across the 24-hours. Cosinor analysis was done with least squares to fit a sine wave to data points in a time series. Hypothesizing that the mice have a daily sleep rhythm, data was aligned to a 24-hour time frame for each mouse across all days, the SWA was normalized by dividing the delta band with all other bands followed by a Hampel filter (outlier detection and method for time series data that uses a sliding window to identify and replace outliers based on the median absolute deviation).

For rhythmicity analysis, a linear mixed-effects (LME) cosinor model was fitted across all valid observations using MATLAB’s fitlmematrix function. The model included fixed effects for MESOR (mean level), cosine, and sine components. The 24-h rhythm was modeled as:

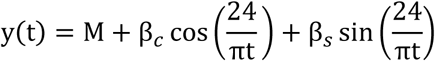

where M is the MESOR, and the amplitude A and acrophase ϕ were derived as:

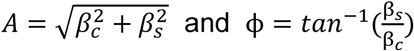

### Sleep homeostasis measures

SH was measured with SWA, defined as power in the delta band in NREM sleep that was normalized to all other frequency bands (theta, alpha, beta and sigma). Decline of SWA over light period was computed with mean SWA values in 10-minute bins over the light period for mice in each group and grouped by week and treatment. To examine how the duration of wakefulness preceding NREM sleep influences subsequent SWA, we analyzed all NREM sleep clusters in each experimental group, defined as union of adjacent NREM bouts that are separated by less than 10 min of non-NREM (wake and REM) and that (i) contain more than 10 minutes of cumulative NREM and (ii) are immediately preceded by more than 10 minutes of continuous wake. The SWA value was taken from the 10th epoch of the NREM cluster to capture stable early-cluster dynamics and reduce effects of brief stage transitions.

### Statistical analysis

Baseline differences in sleep parameters and SWA between groups were analyzed using two-way ANOVA. Weekly seizure distributions were compared using the Mann–Whitney U test, while seizure durations were analyzed using unpaired *t*-tests. One-way ANOVA with appropriate post-hoc corrections for multiple comparisons was employed to compare group differences in interictal spike counts, and sleep measures including SWA between saline or KA-treated groups, as well as KA-treated animals with and without seizures. One-way ANOVA with repeated measures and post-hoc corrections was employed to examine longitudinal changes in sleep and SWA over the 8-week recording period.

Group-averaged power spectral densities (PSDs) were plotted with 95% confidence intervals derived from the *t*-distribution. For each 0.25 Hz frequency bin, a one-way ANOVA across groups was performed, and the resulting *p*-values were corrected for multiple comparisons using the Benjamini–Hochberg false discovery rate (FDR, α = 0.05). Post-hoc Tukey–Kramer pairwise comparisons were used to identify specific group differences.

Cosinor analysis was used to evaluate 24-hour circadian rhythmicity. Between-group comparisons of rhythm parameters were conducted using pairwise contrast matrices applied to the fixed-effects covariance structure of a linear mixed-effects model. Group differences in MESOR, amplitude, and acrophase were tested using the delta method (Milkulich et al, 2003; Hou et al, 2021, Masuda et al, 2023). Mean ± SEM values for SWA and sleep stage transitions were plotted with superimposed cosinor fits to visualize circadian rhythms.

For sleep homeostasis measures, including the decay of SWA during the lights-on period and the increase in SWA following wake, data were pooled by experimental group and analyzed using linear regression and analysis of covariance (ANCOVA). A significant Hour × Group interaction indicated differences in SWA decay or buildup across groups. To quantify week-to-week changes in SWA dynamics, a linear regression model was fitted to SWA during the lights-on period for each week, producing a SWA/hour slope. A second linear regression was then applied to the set of weekly slopes over the 8-week period, and 95% confidence intervals and coefficients of determination (R²) were calculated.

All statistical analyses were performed using MATLAB (MathWorks, Natick, MA) or GraphPad Prism (GraphPad Software, San Diego, CA). Statistical significance was defined as *p* < 0.05.

## Results

### Sleep patterns at baseline in C57 and DBA mice

Analysis of NREM sleep using two-way ANOVA revealed no significant interaction between time and mouse strain [F(11, 36) = 1.40; *p* = 0.3251], but a main effect of time was also observed [F(11, 36) = 11.96; *p* < 0.0001]. Post-hoc Tukey-Kramer multiple comparisons indicated that C57s spent significantly more time in NREM sleep during the first 2-hour block (*p* = 0.001). For REM sleep, two-way ANOVA showed neither a significant interaction between time and mouse strain [F(11, 36) = 1.24; *p* = 0.29] nor a main effect of mouse strain [F(1, 36) = 1.987; *p* = 0.16] (Figure 2A, B). Regarding SWA, two-way ANOVA revealed no significant interaction between time and mouse strain [F(21, 130) = 0.94; *p* = 0.54], but significant main effects of both time [F(21, 130) = 2.52; *p* = 0.008] and mouse strain [F(1, 130) = 327.7; *p* < 0.0001]. Post-hoc Tukey-Kramer multiple comparisons showed that SWA was significantly lower in DBA mice at multiple time points during both lights-on and lights-off periods (*p* < 0.01) (Figure 2C). Data suggest that at baseline, even prior to KA, DBAs have reduced SWA which may be a feature low homeostatic drive.

**Figure 2:**
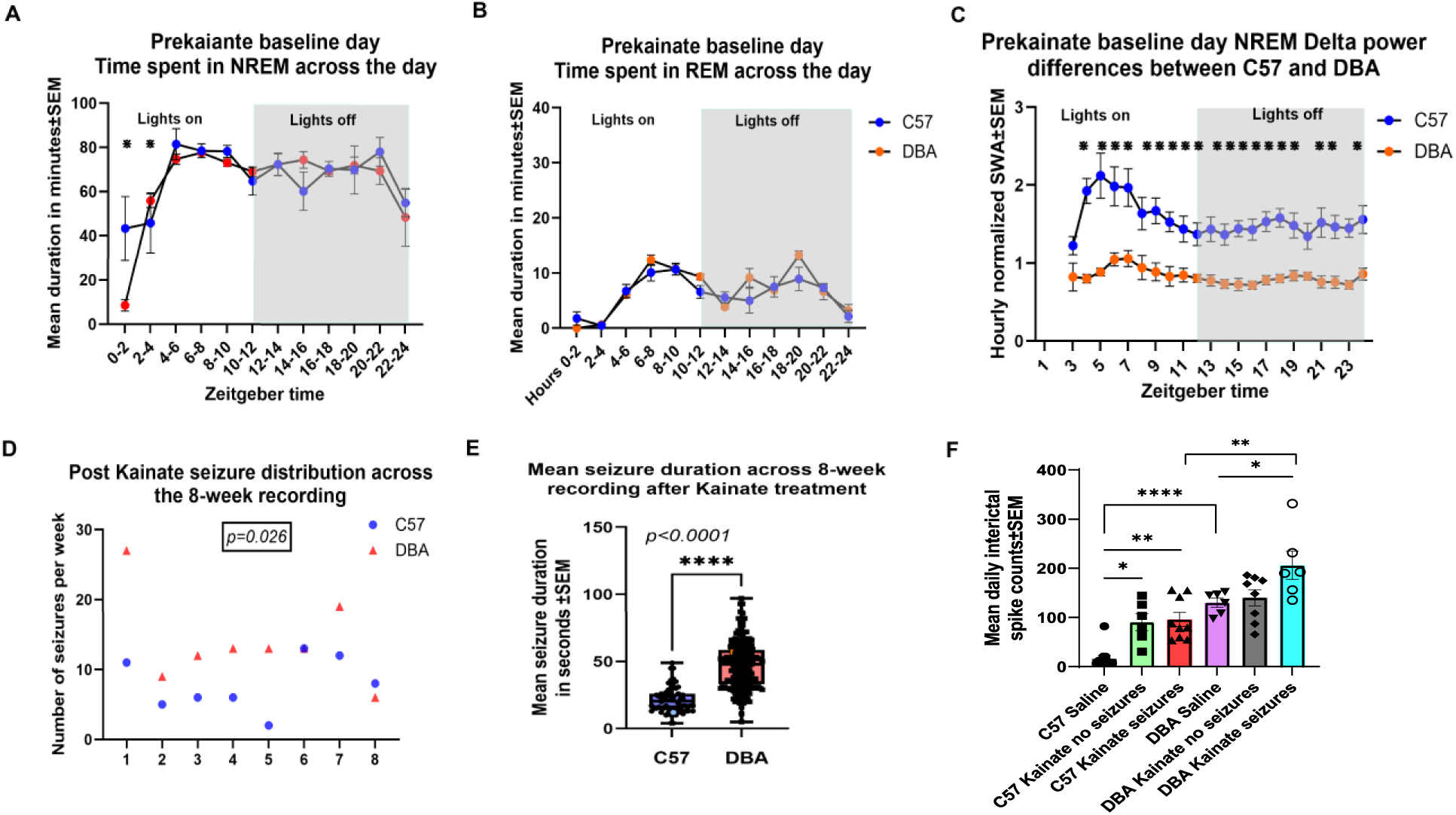
Differences in time spent in NREM (A), REM (B) sleep and SWA (C) are shown in 2-hour blocks for C57 and DBA mice at baseline prior to KA treatment. Differences in each 2-hour block are shown with ←. D: Weekly seizure counts in DBAs than C57s. E: Mean seizure duration on KA-treated C57s and DBAs across the 8-week recording is shown with every seizure shown as a point in the plot E: Group differences in diurnal distribution of seizures are shown for C57s and DBAs (*p<0.05; ** p<0.01; **** p<0.0001).

### Seizures and interictal spikes in KA-treated C57s and DBAs

Seizures were seen as early as week-1 in both groups with no differences in the latency. However, the weekly seizure frequency was higher (Mann-Whitney U-test; p=0.26) (Figure 2D), and seizure duration was longer in DBAs than in C57s (two-tailed t-test; p<0.0001; Figure 2E). Previous reports suggest that interictal spikes can be seen in various strains even without an epileptogenic insult (Pfammater et al, 2018; Purtell et al, 2018; Konduru et al, 2021). Similar to what was suggested in these reports, we detected interictal spikes in all groups including saline-treated groups. There were significant group differences between saline and KA-treated C57s and DBAs (One-way ANOVA- [F (5,40) = 18.36; p<0.0001]). Post-hoc Tukey’s multiple comparison showed that in the saline-treated groups DBAs had greater daily interictal spike frequency than saline-treated than C57s (p<0.0001). Among C57s, KA-treated group without (p=0.015) and with seizures (p=0.001) had significantly more interictal spikes than saline-treated groups. There was no difference in interictal spike frequency between KA-treated C57s with and without seizures (p=0.99). Among DBAs on the other hand, KA-treated group with seizures had greater interictal spike frequency compared to saline-treated group (p=0.035) with no differences in KA-treated animals with and without seizures (p=0.062) (Figure 2F). Finally, KA-treated DBAs had higher daily interictal spike frequency than KA-treated C57s (p=0.0002) (Figure 2F). These data suggest that seizures are more frequent and longer in DBAs. Moreover, interictal spikes are seen in some saline-treated animals though higher in DBAs but increase post KA in both strains regardless of seizures.

### Differences in sleep patterns in saline and KA treated groups

Percent time spent in NREM and REM sleep during lights-on and lights-off periods was compared across all saline- and KA-treated groups. One-way ANOVA showed that were group differences in both strains for NREM and REM: C57s NREM [F (3, 51) = 113.14, *p* < 0.0001]; C57s REM: [F (3, 51) = 143.63, *p* < 0.0001]; DBA mice: NREM: [ F (3, 41) = 81.01, *p* < 0.0001]; REM: [F (3, 41) = 116.13, *p* < 0.0001]. Tukey-Kramer post-hoc analyses revealed that both strains spent significantly more time in sleep during lights-on compared to lights-off (*p* < 0.0001). However, among C57s percent time spent in NREM was not different between KA and saline-treated groups during lights-on (57.59±0.54 vs 59.25±0.45, p>0.05) and lights-off (35.65±1.34 VS 35.134±1.68, p>0.05) (Figure 2B). Time spent in REM was not different between KA and saline-treated C57s during lights on (7.07±0.25 vs 6.59±0.17; p>0.05) or lights off (3.44±0.13 vs 2.97±0.14; p>0.05) (Figure 3A). Similarly in DBAs, time spent in NREM was no different between KA and saline-treated groups during light-on (56.09±1.64 vs 60.29±1.66, p>0.05) or lights-off (37.29±0.97 vs 37.32±1.36, p>0.05) (Figure 3E). REM sleep however was significantly lower in KA-treated DBAs compared to saline both during lights-on (2.32±0.23 vs 5.54±0.16, p<0.0001) and lights-off (1.02±0.14 vs 3.44±0.16; p<0.0001). No significant differences were observed in SWA between saline and KA-treated C57 mice between lights-on and lights-off (*p* > 0.05) (Figure 3C). In contrast, compared to saline-treated ones, KA-treated DBAs showed a significant increase in SWA in both lights-on (0.91±0.12 vs 1.28±0.13, p<0.0001) to lights-off (1.04±0.17 vs 1.40±0.12, *p* < 0.0001) (Figure 3F). The data indicate that KA-treatment disrupts REM in DBAs not in C57s and NREM was unaltered by KA treatment.

**Figure 3:**
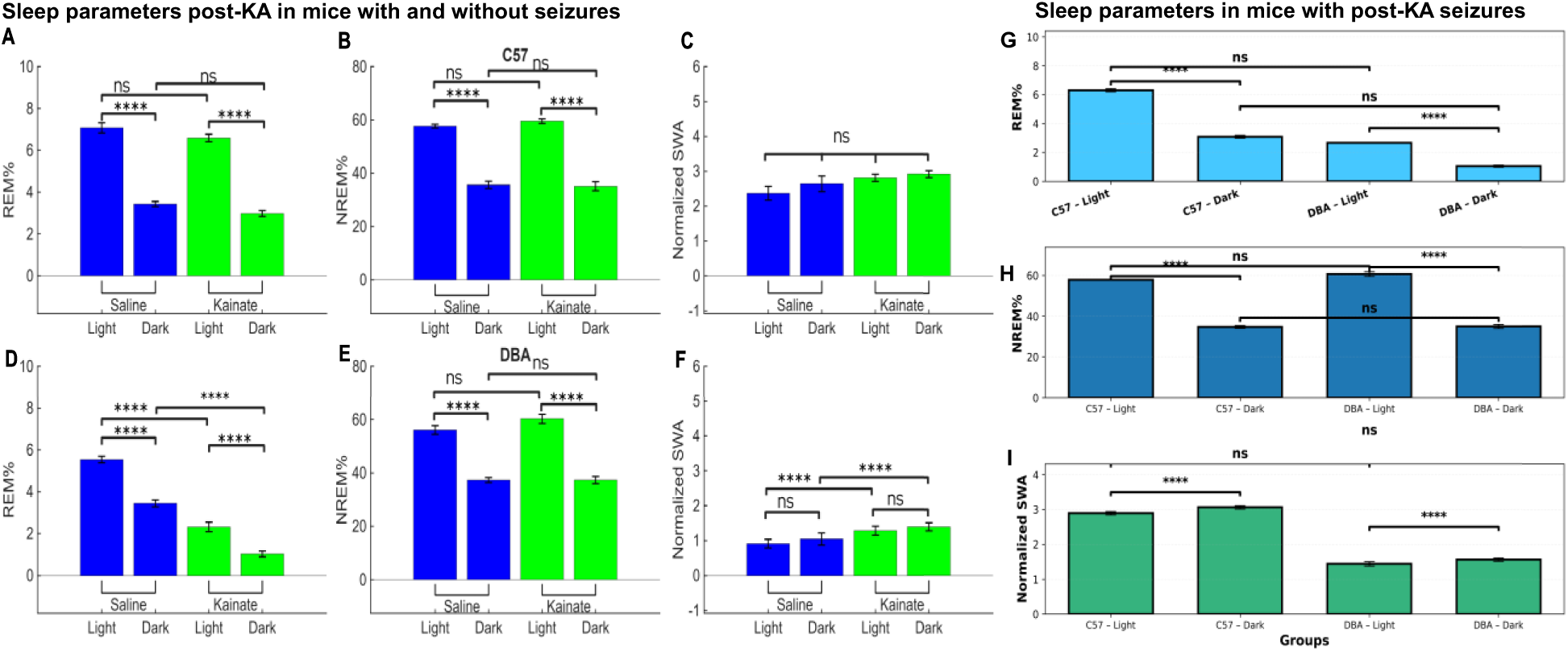
A-F: Differences in percent time spent in NREM, REM, differences in SWA during light-on (lights) and lights-off (dark) in saline and KA-treated C57 and DBAs. A-C: Differences in % time in REM and NREM, and SWA during lights-on and lights-off in saline and KA-treated C57s. D-F: Differences in % time spent in REM and NREM, and SWA during lights-on and lights-off in saline and KA-treated DBAs. G-I: Differences in %time spent in REM and NREM sleep, and SWA in C57s and DBAs that developed post-KA spontaneous seizures lights-on (light) and lights-off (dark) (**** = p<0.0001).

Analysis restricted to KA-treated animals that developed seizures only showed group differences in percent time in NREM [F (3, 29) = 187.76, *p* < 0.0001], REM [F (3, 29) = 348.58, *p* < 0.0001] and in SWA [F (3, 29) = 33.50, *p* < 0.0001]. Post-hoc multiple comparisons revealed that compared to C57s, DBAs had significantly less percent time in REM sleep during both lights-on (6.30±0.11 vs 2.66±0.13, p<0.0001) and lights-off (3.08±0.10 vs 1.05±0.08, -<0.0001) (Figure 3G). On the other hand, percent time NREM sleep was significantly less in C57s that DBAs in lights-on (57.85±0.53 vs 60.69±1.14, p<0.0001) and lights off (34.73±0.71 vs 34.95±1.01, p<0.0001) (Figure 3H). Mean SWA was significantly lower in DBA mice compared to C57s during both lights-on (1.44±0.06 vs 2.89±0.05; p<0.0001) and lights-off (1.56±0.05 vs 3.06±0.05, p<0.0001) (Figure 3I). Together, these data indicate that strain-specific sleep disruptions in KA-treated groups with seizures, where REM sleep and SWA are particularly vulnerable to disruption in the seizure-susceptible DBAs.

Sleep fragmentation was assessed using sleep bout durations in all saline and KA-treated groups. There were differences seen in both NREM and REM in both C57s and DBAs (One-way ANOVA-C57s: [F (3, 51) = 174.10; p<0.0001]; DBAs: [F (3, 41) = 124.56; p<0.0001]). Post-hoc multiple comparisons showed that in all saline and KA-treated groups bout durations for NREM or REM were greater in lights-on compared to lights off (p>0.0001) (Figure S1 A-D). However, no difference in bout duration was both in NREM and REM between KA and saline C57s during light-on or lights-off (p>0.05) (Figure S1 A&B). DBAs on the other hand and increase in NREM bout length in lights-off compared to lights-on (6.68±0.21 min vs 6.23±0.18 min; p=0.03) (Figure S1 C). Additionally, REM bout durations were lower in KA-treated DBAs both in lights-on (0.77±0.05) and lights-off (0.38±0.03) compared to saline group (1.26±0.03 and 0.90±0.04 min respectively; p<0.0001) (Figure S1 D). Overall, REM sleep was much more fragmented in KA-treated DBAs.

### Changes in sleep patterns across 8-week recording through epileptogenesis

To assess whether sleep patterns progressively deteriorate in animals that developed spontaneous seizures following KA, we examined weekly changes in percent time spent in NREM and REM sleep, as well as slow wave activity (SWA), over the 8-week recording period. Among C57s that developed seizures, there was no significant change in the percent time spent in NREM sleep from week 1 to week 8 [One-way ANOVA with repeated measures: F (7, 179) = 0.697; *p* = 0.67] (Figure S2-A). In contrast, DBA mice showed a significant week-to-week difference in NREM sleep time [F(7, 179)= 4.87, p=0.03], with post-hoc comparisons revealing increased NREM sleep in week 6 (*p* = 0.005), week 7 (*p* = 0.04), and week 8 (*p* = 0.01) compared to week 1 (Figure S2-A). No significant differences in percent time spent in REM sleep were observed across the 8 weeks in either C57 (Figure S2-C) or DBA mice (Figure S3-B) (F (7, 179) = 0.355; p> 0.05). Analysis of SWA revealed a significant effect of time in C57 mice [F (7, 179) = 2.355; *p*= 0.025], with post-hoc tests indicating reduced SWA in week 8 compared to week 2 (*p* = 0.037) (Figure S2-C). In DBA mice, no significant changes in NREM SWA were detected across the 8-week period (*p* > 0.05) (Figure S2-C). Overall, there was no consistent or progressive deterioration in sleep patterns across the epileptogenesis period. While C57 mice exhibited a late decrease in SWA, and DBA mice showed an increase in NREM sleep time during later weeks.

### Power spectral density

To further investigate the sleep differences between C57 and DBA mice with seizures—particularly differences in SWA—we performed power spectral density (PSD) analysis during NREM sleep. PSD was calculated from frontal EEG recordings during NREM sleep across the entire 24-hour period for all C57 and DBA mice, including saline-treated and KA-treated animals with and without seizures. Group differences were assessed with and one-way ANOVA and post-hoc Tukey-Kramer tests at every bin. Analyses showed that all C57 groups exhibited significantly higher power in the delta frequency band of 1.25-3.75 Hz compared to all DBA groups (*p* < 0.05). DBAs on the other hand showed significantly higher power across theta, alpha, and beta bands (5–14 Hz) relative to C57s (*p* < 0.05). C57s demonstrated greater power than DBA mice in low gamma band (17-20 Hz) (*p* < 0.05) (Figure 4). These findings highlight consistent spectral differences in NREM sleep between strains, with C57 mice showing enhanced low-frequency (delta) power and DBA mice showing elevated mid-frequency (theta to beta) activity. Notably, these differences persisted regardless of treatment condition (saline or KA) or seizure status. Only statistically significant post-hoc differences between all C57 and all DBA groups in each frequency bin were reported here.

**Figure 4:**
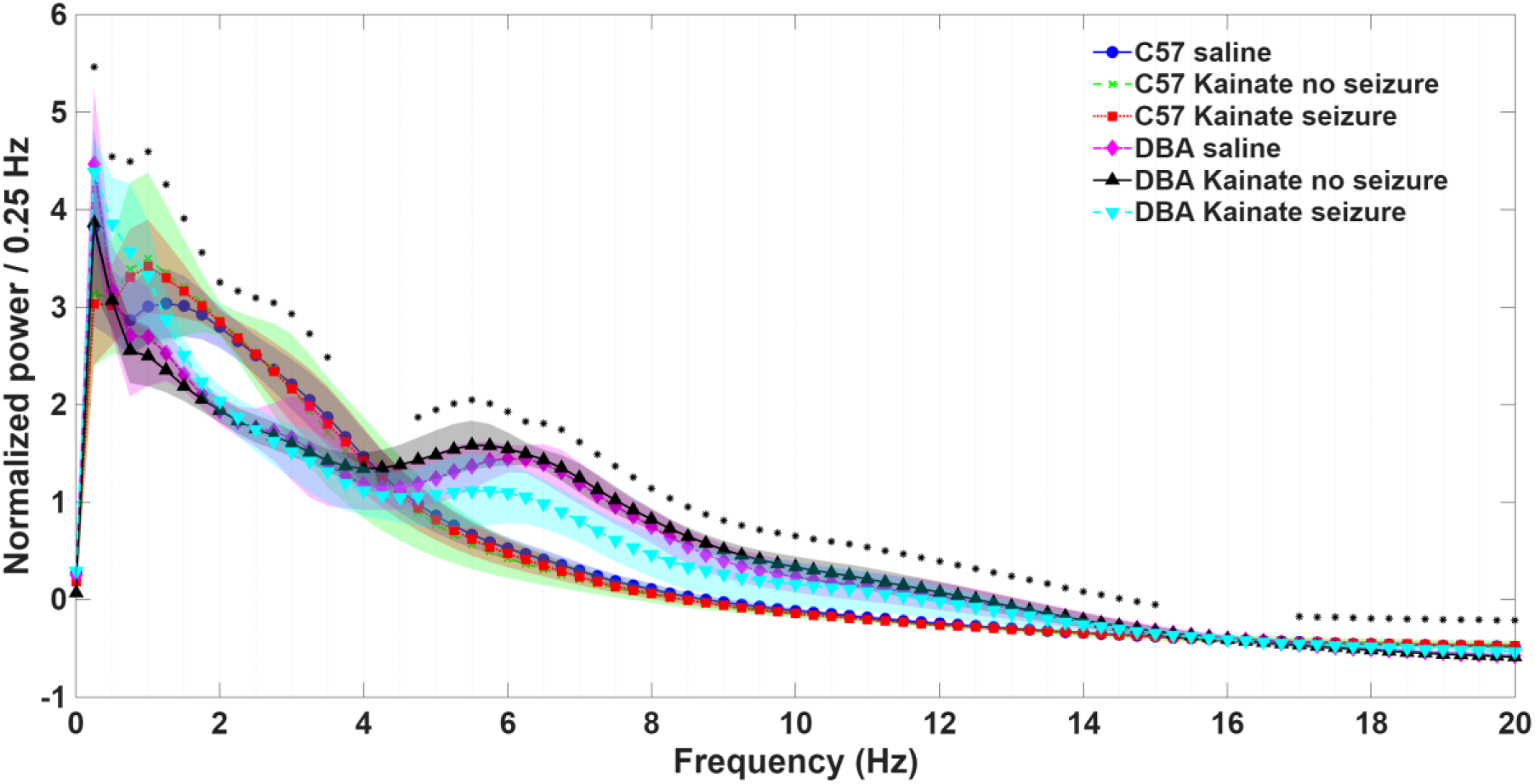
Differences in power spectral density is shown for C57s and DBAs treated with saline, treated with KA but with and without seizures. Data shows that all C57s with KA-seizures followed by KA+ seizures and C57s with saline had higher power in delta band (0.5-4hz) whereas DBAs all groups had higher power in the theta, alpha and beta bands (5-14 Hz). Differences between groups at every 0.25Hz frequency band is shown with a * indicating p<0.05). The means for each group along with 95% Confidence intervals are shown.

### Circadian rhythm of SWA in C57s and DBAs with seizures

Given the observed differences in sleep patterns and SWA between C57s and DBAs with seizures, we next investigated whether the periodicity of circadian rhythms changed from before to after onset of spontaneous seizures. Cosinor analysis revealed that both C57 and DBA mice exhibited significant diurnal rhythms in SWA though the overall amplitude of the circadian rhythm was higher in C57s than in DBAs (Figure 5A; Table 1). In the linear mixed-effects cosinor model, each group’s rhythmic component was represented by two fixed-effects coefficients: the cosine term and the sine term which jointly define the amplitude and phase of the 24-hour oscillation. Data showed that rhythmicity, MESOR, acrophase and amplitude of SWA were all different between C57s before and after seizures whereas no such differences were seen in DBAs (Table S1). In addition, we performed Cosinor analysis to examine the circadian changes in transitions in sleep stages from wake to NREM or REM, from NREM to wake or REM and from REM to NREM or wake (Figure 5B-G). Data showed that in C57s there was a significant difference in transitions from NREM to REM and REM to NREM but not in transitions between other sleep stages from before and after onset of seizures. In DBAs on the other hand, no differences were seen in any stage transitions (Table 2). These data suggest that circadian rhythms in C57s likely exhibit plasticity and adapt to physiologic stressors whereas DBAs on the other hand have impaired adaptability.

**Figure 5:**
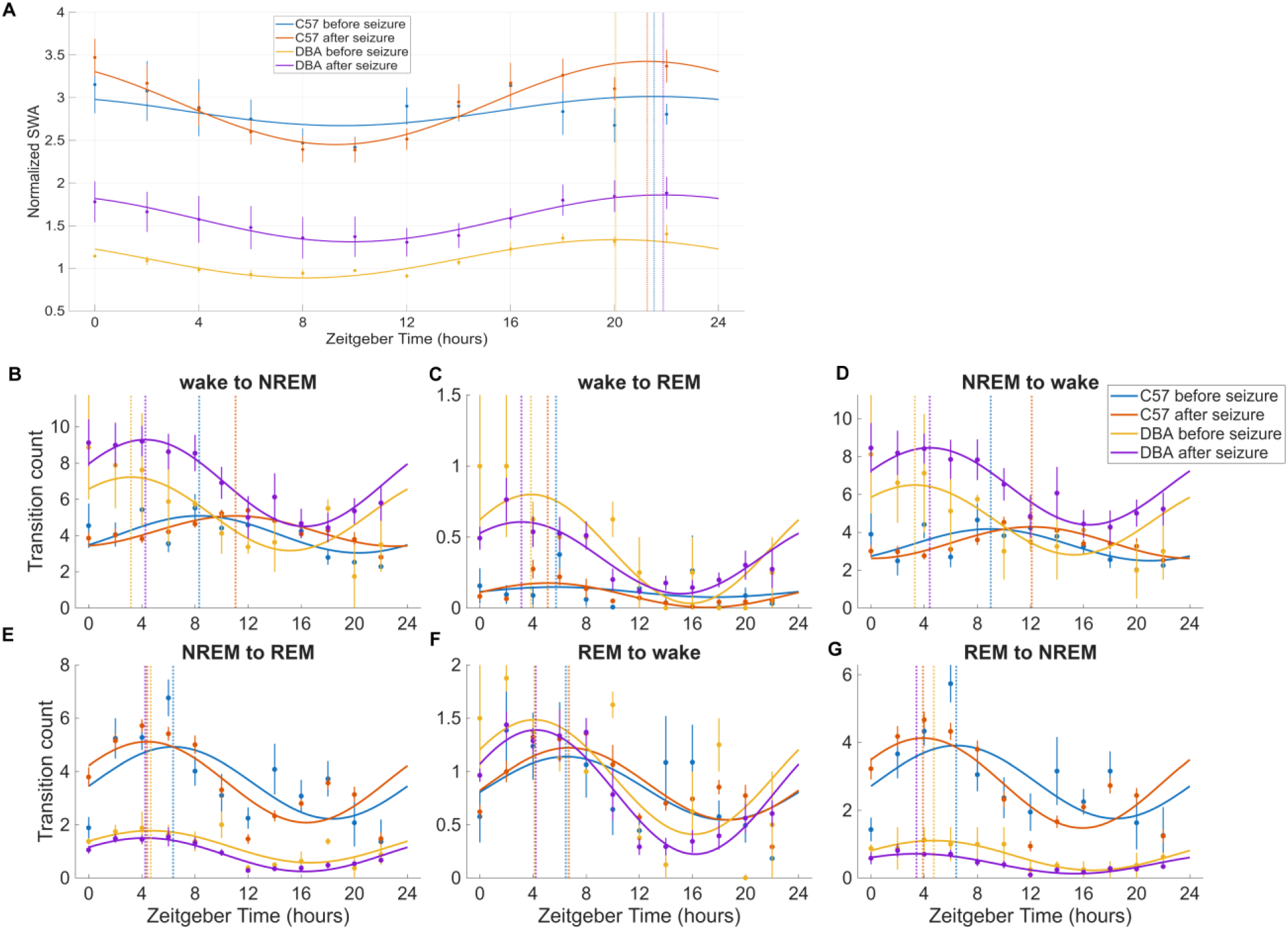
A: Differences in diurnal rhythm of NREM slow-wave activity (SWA) across the day before vs. after the first seizure in C57 and DBA mice assessed by Cosinor analysis: A: Diurnal rhythm of SWA is shown. Points show group mean ± SEM at bin centers; colored solid lines overlay one-component 24-h cosinor fits evaluated on a dense time grid. Vertical lines represent the Acrophase for each group. B-G: Diurnal rhythms of vigilance-state transitions: For C57s and DBAs treated with KA, we computed counts for six state transitions per bin: wake to NREM (B), wake to REM (C), NREM to wake (D), NREM to REM (E), REM to wake (F), REM to NREM (G), before and after they developed spontaneous seizures. Vertical lines show the fitted acrophase (peak time) for each group in each transition panel. Statistical data shown in table 2.

**Table 1:**
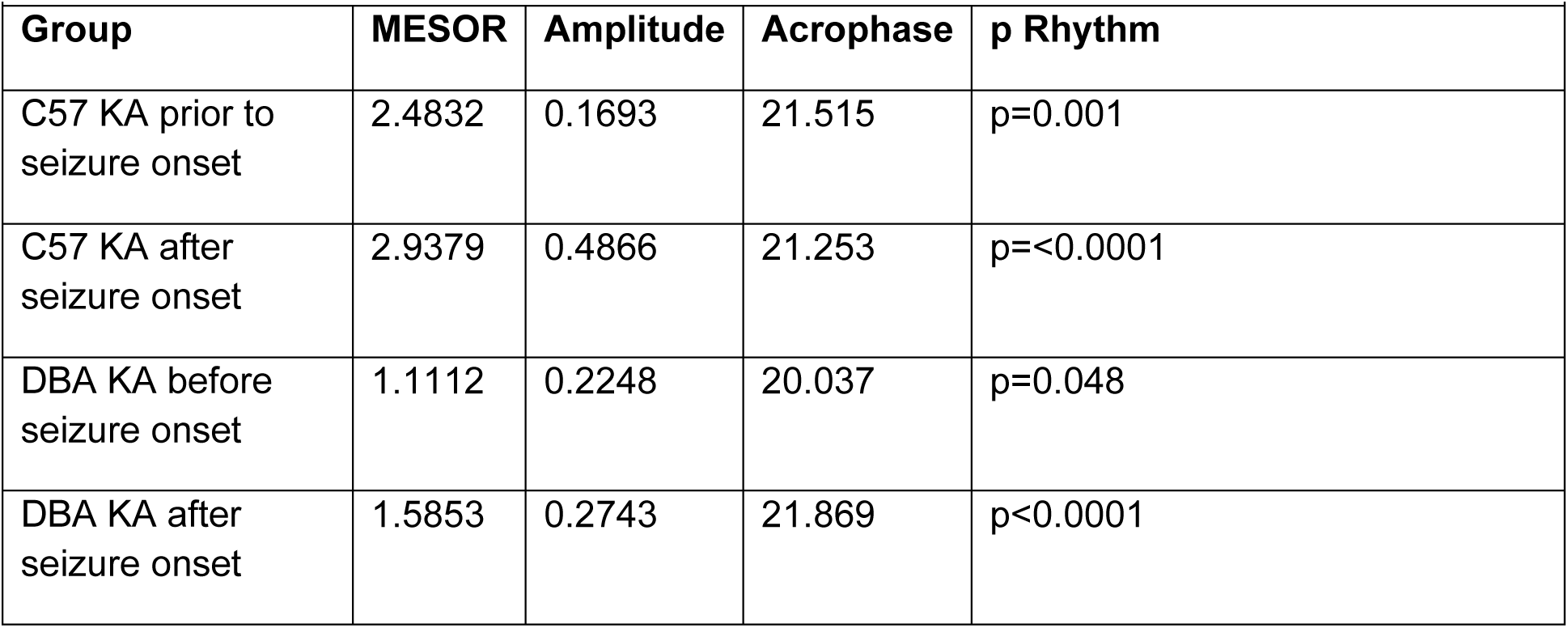
Results of Cosinor analysis of slow wave activity rhythmicity and group differences:

| Group | MESOR | Amplitude | Acrophase | p Rhythm |
| --- | --- | --- | --- | --- |
| C57 KA prior to seizure onset | 2.4832 | 0.1693 | 21.515 | p=0.001 |
| C57 KA after seizure onset | 2.9379 | 0.4866 | 21.253 | p=<0.0001 |
| DBA KA before seizure onset | 1.1112 | 0.2248 | 20.037 | p=0.048 |
| DBA KA after seizure onset | 1.5853 | 0.2743 | 21.869 | p<0.0001 |

**Table 2:**
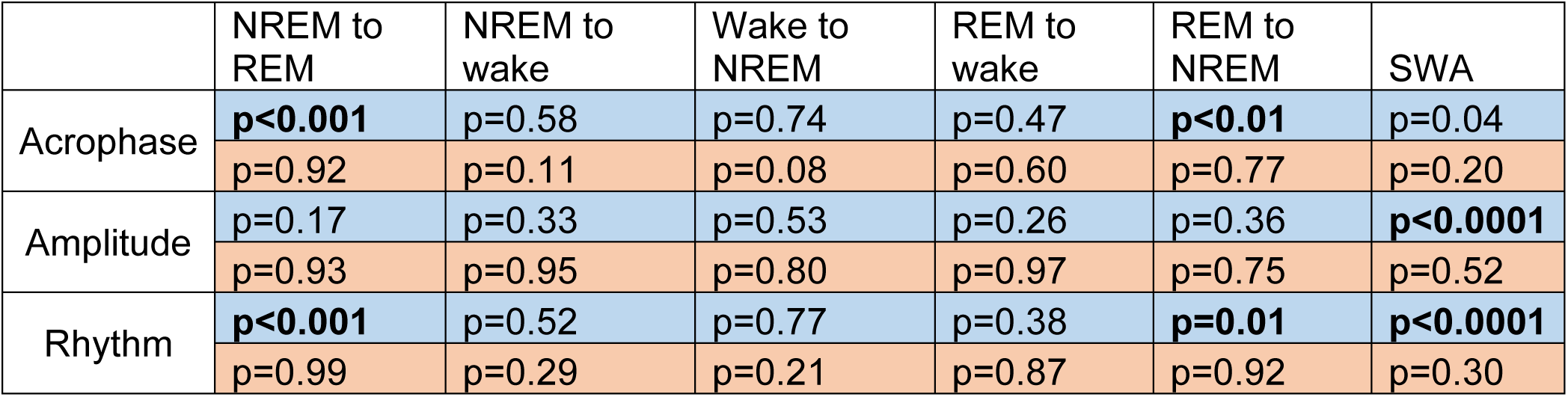
Statistical data analysis for the cosinor analysis of stage transitions is shown with MESOR, Amplitude, Acrophase, R2 for the fit and F Rhythm (C57s before and after onset of seizures in blue and DBAs before and after onset of seizures in orange).

### Sleep homeostasis measures in C57 and DBA mice

Sleep homeostasis measures including decay of SWA (slope per minute) across lights-on as mice sleep and rise in SWA with time spent awake (after sleep onset) was examined using linear regression models. When data was plotted for all groups and all days, the SWA decayed from lights-on to lights-off in all groups though the Y-intercepts were different. The slopes for saline-treated C57s, KA-treated C57s without seizures and KA-treated C57s with seizures respectively were -0.070 (95%CI: -0.079 to - 0.061), -0.095 (95% CI: -0.107 to -0.082) and -0.09 (95% CI: -0.10 to -0.082). The slopes for saline-treated and KA-treated DBAs with and without seizures respectively were -0.030 (95% CI: -0.043 to 0.017), -0.035 (95%CI: -0.046 to -0.024) and -0.051 (95%CI: -0.064 to -0.038) (Figure 6A). ANCOVA analysis showed that the slopes were significantly different [F (11, 3302) = 729.4; p<0.0001] with post-hoc pairwise comparisons showing all C57 groups were different from each other (p<0.05) whereas among DBAs only difference in slopes was between saline-treated and KA-treated DBAs with seizures (p<0.05) (Table 3). These data suggest that homeostatic drive measured by decay of SWA during lights-on was preserved (negative values for all groups) but with some differences between groups and as seen in Figure 6A, the Y-intercepts were lower for DBAs.

**Figure 6:**
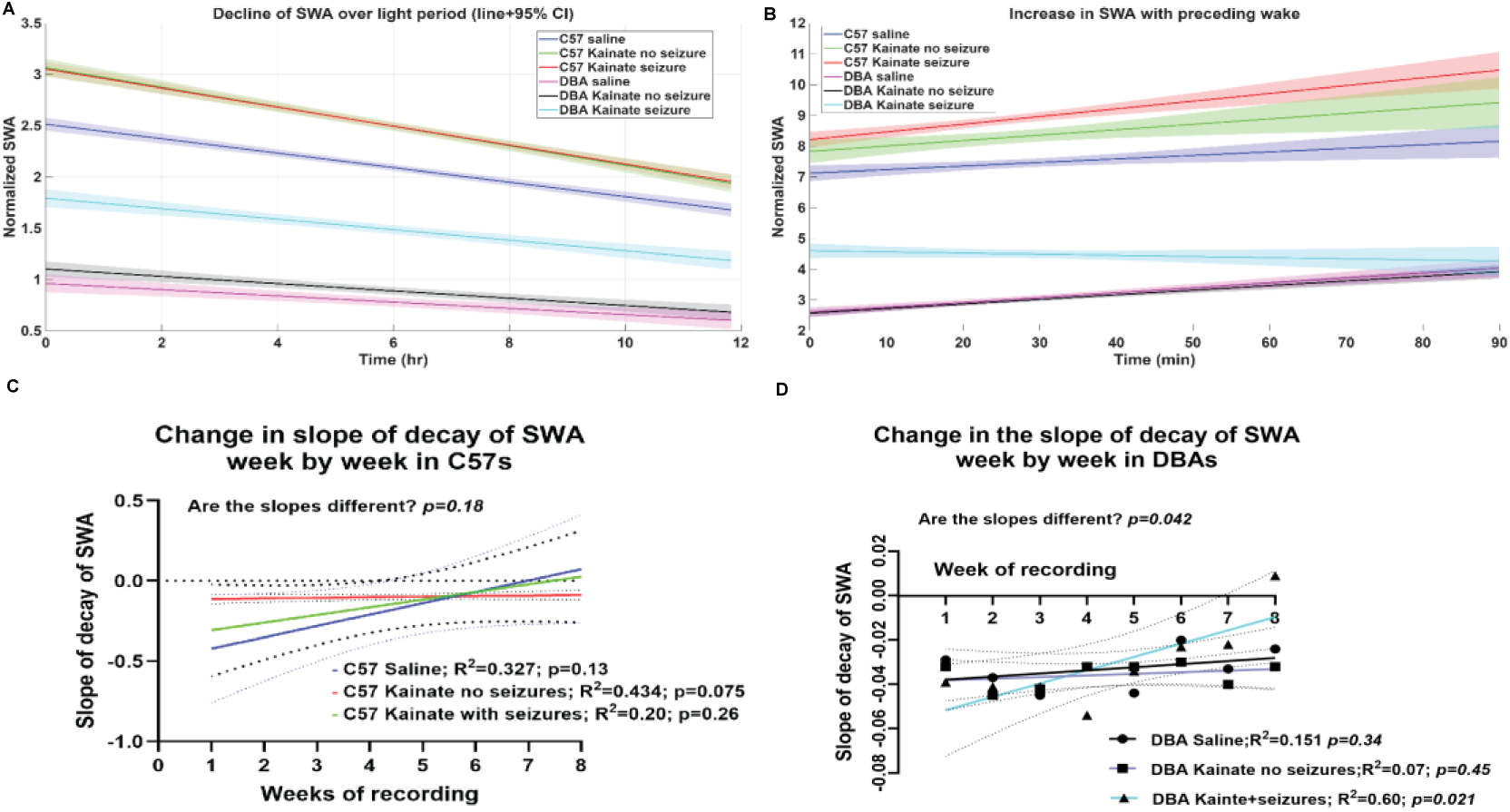
Measures of sleep homeostasis in C57 and DBAs that are saline-treated and KA-treated with or without seizures. A: decay of SWA from the beginning of lights-on to end of lights-on in for C57s and DBAs treated with saline, KA with and KA without seizures. B: Rise in SWA with preceding wake after sleep onset is shown all C57 and DBA groups. C and D: Change in the slopes of decay of SWA during lights off from week 1 to week 8 in each of the treatment groups are shown fitted with a linear regression.

**Table 3:** Differences in overnight decay of SWA and rise with preceding wake between groups.

| Treatment group | Decline in SWA over light period |  | Rise in SWA with Preceding wake |  |
| --- | --- | --- | --- | --- |
|  | Difference in slope per min | p value | Difference in slope per min | p value |
| C57 saline vs C57 KA no seizure | 0.024 | **0.002** | -0.007 | 0.17 |
| C57 saline vs C57 KA seizures | 0.022 | **0.002** | -0.013 | **0.003** |
| C57 KA + vs - seizures | -0.002 | 0.75 | -0.005 | 0.34 |
| DBA saline vs<br>DBA KA no<br>seizures | 0.005 | 0.54 | 0.001 | 0.76 |
| DBA saline vs<br>DBA KA<br>seizures | 0.02 | **0.024** | 0.018 | **0.008** |
| DBA KA + vs -<br>seizures | 0.015 | 0.07 | 0.016 | **0.001** |

With regard to the rise in sleep pressure with preceding wake, KA treated C57s with seizures had the steepest rising slope (0.030; 95% CI 0.023-0.037) compared to KA treated C57s without seizures (0.024; 95% CI 0.015-0.034) or saline-treated group (0.016; 95% CI 0.010-0.022). Saline-treated DBAs (0.016; 95% CI 0.01-0.023) and KA-treated DBAs without seizures (0.015; 95%CI 0.009-0.021) had upgoing slopes as well, indicative of rising sleep pressure with preceding wake. However, KA-treated DBA with seizures had a down going slope (-0.001; 09%CI -0.009 to -0.006) indictive of loss of homeostatic increase in SWA. Differences in the slopes were significant with ANCOVA [F (11, 9191=843.6; p<0.0001). The Y-intercepts were lower for all DBA groups compared to C57 groups (Figure 6B). Post-hoc Bonferroni corrections (within C57s and DBAs) showed that C57 saline group was different from KA-treated group with seizures (p=0.003) and no difference seen in C57 KA group with and without seizures (p=0.34). Among DBAs, saline-treated group was different from KA group with seizures (p=0.0008) and KA-treated groups with and without seizures were also different from each other (p=0.001). These data suggest that the SH measured by rise of SWA with preceding wake is relatively preserved in C57s regardless of KA treatment and in saline or KA-treated DBAs without seizures, but seizures in DBA after KA resulted in an impairment of this SH measure.

Lastly, we attempted to examine if there is a progressive change in sleep homeostatic drive in C57s and DBAs treated with saline and KA groups with and without seizures on a week-to-week basis across the 8-week recording. Using a linear regression of the fitted slopes of decay in each week, data showed that all C57 groups had a decay of SWA in every week without a clear progressive change from week 1-8 (C57 saline p=0.13; C57 KA without seizures, p=0.07; C57 KA with seizures p=0.26; Difference in the slopes-p=0.18; Figure 6C). DBAs that are saline-treated (p=0.34) and KA-treated DBAs without seizures (p-0.45) also had relatively unchanged slope across the 8-weeks. However, in KA-treated DBAs with seizures, the slope was rising from week 1-8 (p=0.08) with slopes between all DBA groups being different (p=0.042) (Figure 6D) (Table S2). The actual slopes in each week in each treatment group are shown in Figure S2. These data suggest that seizures in C57 do not result in progressive change in SH where KA-treated DBAs with seizures had a progressively change across the 8-week recording.

## Discussion

This study investigated how baseline sleep characteristics and sleep disturbances following KA administration relate to the development of spontaneous seizures in two distinct mouse strains, one considered epilepsy resistant-C57BL/6J (C57) and the other epilepsy susceptible-DBA/2J (DBA). Our data provide evidence that differences in sleep, particularly in SWA prior to KA in DBAs. Following KA, DBAs have worse seizures, more disruptions in sleep, circadian rhythms as well as SH. The disruptions in sleep, circadian rhythms and SH may be factors associated with increased epileptogenesis in DBAs compared to C57s.

### Baseline Differences and Seizure Susceptibility

At baseline, both C57 and DBA mice spent relatively similar amounts of time in NREM and REM sleep, but SWA was significantly lower in DBAs. SWA is a well-established marker of sleep pressure or SH and cortical synchrony (Esser et al, 2007). According to the synaptic homeostasis hypothesis SWA relates to cortical synaptic strength and network synchronization which increases as sleep debt accumulates across wake time and diminishes with sleep (Tononi and Cirelli, 2020). Persistently low SWA (as seen in DBAs) had been linked to disrupted thalamocortical function and weakened synaptic strength (Tononi, 2009). Similar to our observations, one prior study showed reduced SWA at baseline in DBAs compared to C57s (Hasan et al, 2012). The lower SWA in DBAs may reflect a baseline deficit in sleep regulatory mechanisms, reduced sleep pressure and synaptic plasticity. There is no prior work to inform whether baseline deficits in sleep homeostatic drive predisposes to epileptogenesis. Prior work in Drosophila was contradictory where it was the increased but not reduced homeostatic drive that worsened seizures (Cuddapah et al, 2025). DBAs here exhibited reduced sleep pressure, i.e., reduced SWA, yet had more frequent and prolonged seizures than C57s. While further work is needed in support, we hypothesize that the deficits in SWA at baseline in DBAs reflecting reduced homeostatic drive, may be a factor in rendering them more epilepsy susceptible following KA.

### Post-KA Sleep Disruption

There is a dearth of prior studies where sleep was examined with chronic EEG recordings in mouse models of epilepsy. One study showed reduced slow wave and REM sleep post-KA but the recordings were short (Ayala-Guerrero et al, 2002). Reduced REM sleep was also noted in a model of sleep-related hypermotor epilepsy but the recordings were 24-48 hours (Wang et al, 2025). In cats however, KA treatment resulted in reduced NREM as well as REM sleep bouts that persisted for 6-7 weeks (Szumusiak and McGinty, 1986). In a mouse model of severe traumatic brain injury, sleep homeostatic drive was disrupted acutely as well as chronically, where the power in NREM SWA increased after TBI along with a loss of diurnal oscillation (Konduru et al, 2021). In our experiments, DBAs exhibited deficits in percent REM sleep and REM bout duration following KA treatment. Additionally, KA treatment did not result in any changes in NREM in both C57s and DBAs. Interestingly when KA-treated animals that developed seizures were examined, DBAs had reduced REM, increased NREM sleep and lower SWA compared to C57s. This consistent pattern suggests that DBAs have impaired REM regulation and decreased SWA, especially when they develop seizures. We hypothesize that the disruptions in sleep and homeostatic drive following KA may contribute to progression of epileptogenesis in DBAs.

### Temporal Dynamics of Sleep Changes During Epileptogenesis

Several studies have implied the progressive nature of temporal lobe epilepsy in humans and in the KA model in rodents (Sutula, 2004; Bernhardt, 2009). While sleep and epilepsy have a bidirectional relationship (Maganti and Jones, 2021), no studies to date have examine how sleep patterns evolve progressively through epileptogenesis with chronic recordings. We found only subtle changes in sleep architecture over the 8-week recording period where DBAs with seizures did show a late increase in NREM duration (weeks 6–8), while it remained stable in C57s. SWA in declined in C57s in week 8 post-seizure onset, while remaining low and unchanged in DBAs. As seizures disrupt sleep, an increase in SWA over time may be compensatory, due plasticity in SH. A lack of such an increase in DBAs may suggest that this form of homeostatic plasticity is impaired over time during epileptogenesis in the epilepsy-susceptible strain.

### Altered EEG power spectrum and Diurnal Rhythms

Power spectral density (PSD) analysis done previously in a Kainic model showed that C57s that develop seizures had reduced theta and increased gamma band power (Duglazde et al, 2007). Consistent with the initial finding of increased SWA, PSD analysis also showed higher power in the delta band in KA-treated C57s, while DBAs had higher power in the theta, alpha, and beta bands. It is possible that the higher seizure duration and frequency as well as higher interictal activity in DBAs could have contributed to the higher power in theta to beta bands. Prior studies have suggested that the increase in higher frequency bands in NREM sleep may reflect cortical hyperexcitability (Dunstan et al, 2025) and impaired sleep regulation (Halasz et al, 2023), both of which are implicated in epileptogenesis.

In the cosinor model, a cosine curve is fitted to periodic data within a regression model and is a commonly used method to describe cyclical variation (Klerman et al, 2016). There are no prior studies to our knowledge that analyzed circadian rhythm of sleep using Cosinor analysis in mouse models of epilepsy. We show that both strains maintained a significant diurnal rhythm in SWA before and after seizure onset, though the rhythm is strengthened (higher amplitude) in C57s and shifted acrophase after onset of spontaneous seizures. In contrast, the circadian rhythm and acrophase did not change after onset of spontaneous seizures along with consistent low SWA amplitudes across the day in DBAs. This suggests impaired circadian regulation of sleep in DBAs, we hypothesize that this contributes to acceleration of epileptogenesis. Furthermore, diurnal patterns of vigilance state transitions were altered in both strains, with C57s showing significant acrophase shifts post-seizure. DBAs did not exhibit this rhythmic adaptation. Overall, these data suggest that DBAs may have a less flexible circadian timing system that is not amenable to perturbations in disease states such as epilepsy.

### Sleep Homeostasis and Its Disruption in Epilepsy

A limited number of studies examined SH in epilepsy. In patients with epilepsy, increased SWA power was observed and overnight decay of slope of slow waves was lost indicative of disruptive homeostasis (Boly et al, 2017). Interestingly patients who succumbed to SUDEP, who generally have more severe seizures and mortality during sleep, SH was disrupted with no decay of SWA across sleep, similar to what was seen in DBAs with seizures after KA (Magana-Tellez et al, 2025). C57 mice exhibited the expected decay in SWA across the lights-on period and a corresponding rise in SWA with preceding wake suggesting that epilepsy resistant mice have intact regulation of SH that may possibly be protective against development of spontaneous seizures. In stark contrast, a key finding was that DBAs with seizures showed a flattened slope of decay of SWA during lights on and no increase in SWA with preceding wake suggestive of disrupted SH. This suggests a fundamental breakdown in the mechanisms that regulate SH in this strain following epileptogenesis. Notably, the homeostatic decay of SWA progressively changed over the 8-weeks in DBAs, further supporting the idea of a link between impaired SH and the maintenance or progression of epilepsy.

### Implications and Future Directions

Collectively, these findings suggest that pre-existing differences in SH and architecture — particularly lower SWA and impaired REM regulation — may potentially be factors that predispose certain mouse strains such as DBA to more severe epileptogenic outcomes following an initial insult such as KA-induced status epilepticus. Furthermore, the inability to maintain circadian rhythms and abnormal SH may be indicative of impaired neuroplasticity as well disrupted thalamocortical regulation, both hallmarks of epilepsy (Halasz et al, 2023). The strong diurnal rhythm and preserved homeostasis seen in C57s, even post-seizure, may reflect protective mechanisms that render them resistant to epileptogenesis.

Understanding these strain-specific differences could guide the development of sleep-based biomarkers or interventions to identify or even prevent epileptogenesis in at-risk individuals.

There are several limitations that should be pointed out. First, there may be strain-specific confounds unrelated to epilepsy (e.g., metabolic or behavioral differences) that resulted in our findings. Not all parameters were analyzed in saline, KA treatment without or with spontaneous seizures. EEG recordings were done 5-days a week and we may have missed seizures on days recordings were not done. Finally, we have not instituted any treatment to suppress seizures or to enhance SH to understand whether the observations have a cause-and-effect relationship.

## Conclusion

In conclusion, this study underscores the importance of sleep and its regulation in the pathophysiology of epilepsy. Pre-existing differences in SWA and REM architecture may influence susceptibility to epilepsy, while impaired SH and disrupted diurnal rhythms following seizure onset may potentially contribute to disease maintenance or progression. The translational significance is that these findings support further investigation into sleep-targeted therapies as an approach to modulate epileptogenesis and improve outcomes in epilepsy.

## Acknowledgement

This work is supported by a CDMRP (Department of War) Grant: PR 221869 (PI: RM)

