## Supplementary material for "Sleep dynamics and epileptogenesis following Kainic acid in epilepsy-susceptible (DBA/2J) and epilepsy-resistant (C57BL/6) mice"

Figure S1: Different in bout duration for NREM and REM sleep were shown. NREM bouts in C57s (A) and DBAs (C) were longer in lights-on than lights-off for saline and KA-treated groups. NREM bout duration remained no different between KA and saline groups in lights-on but increased post KA during lights-off. REM bout durations were also greater in lights-on compared to lights-off in both C57s (B) and DBAs (D). They decreased post-KA both during lights-on and lights off in both groups (*p<0.05; ** p<0.01; **** p<0.0001).

Figure S2: Figure 4: Change in sleep and SWA (± SEM) from week-1 to week-8 are shown. A: Percent time spent in NREM in C57s and DBAs (A). Percent time in REM in C57s and DBAs (B). SWA in C57s and DBAs (C).

Figure S3: A-F show how decay of SWA during lights off changes from weeks 1-8 of the recording in C57 saline (A), C57 KA without seizures (B), C57 KA with seizures (C); DBA saline (D), DBA KA without seizures (E) and DBA KA with seizures (F). Note all groups had a decay of SWA during lights-on from week-1 week-8 excepting KA-treated DBAs with seizures had a loss of decay of SWA in weeks 7-8.

Figure S1:

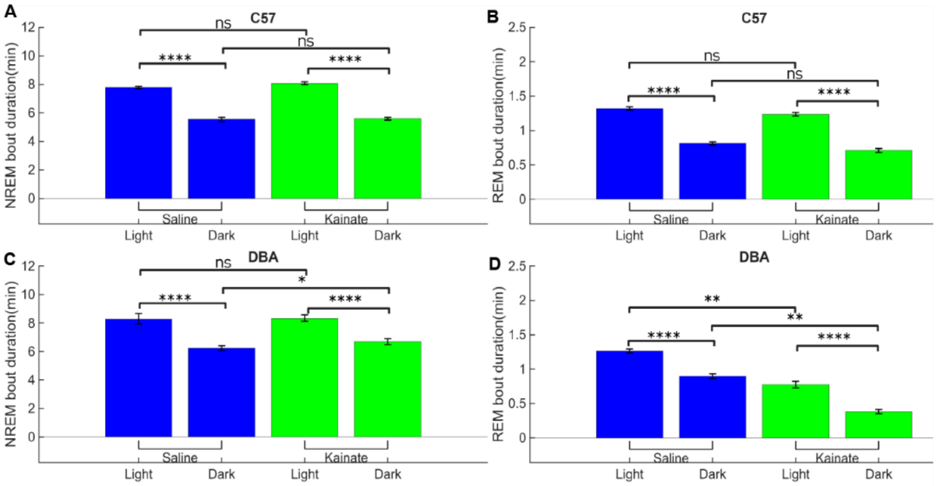


Figure S2:

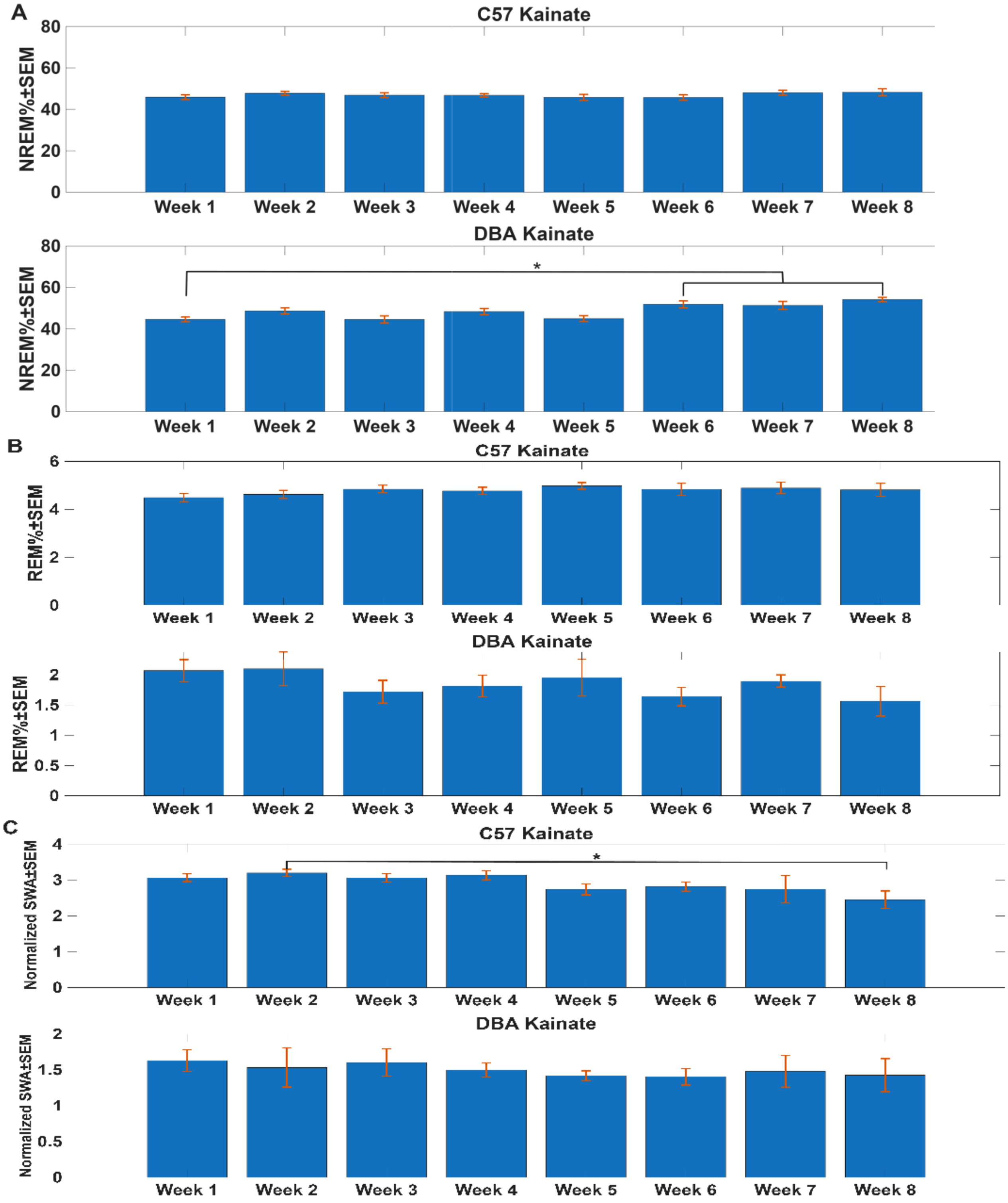
A
C57 Kainate
NREM%±SEM
80
60
40
20
0
Week 1
Week 2
Week 3
Week 4
Week 5
Week 6
Week 7
Week 8
DBA Kainate
*
B
C57 Kainate
REM%±SEM
6
4
2
DBA Kainate
1.5
1
0.5
C
Normalized SWA±SEM
3

Figure S3:

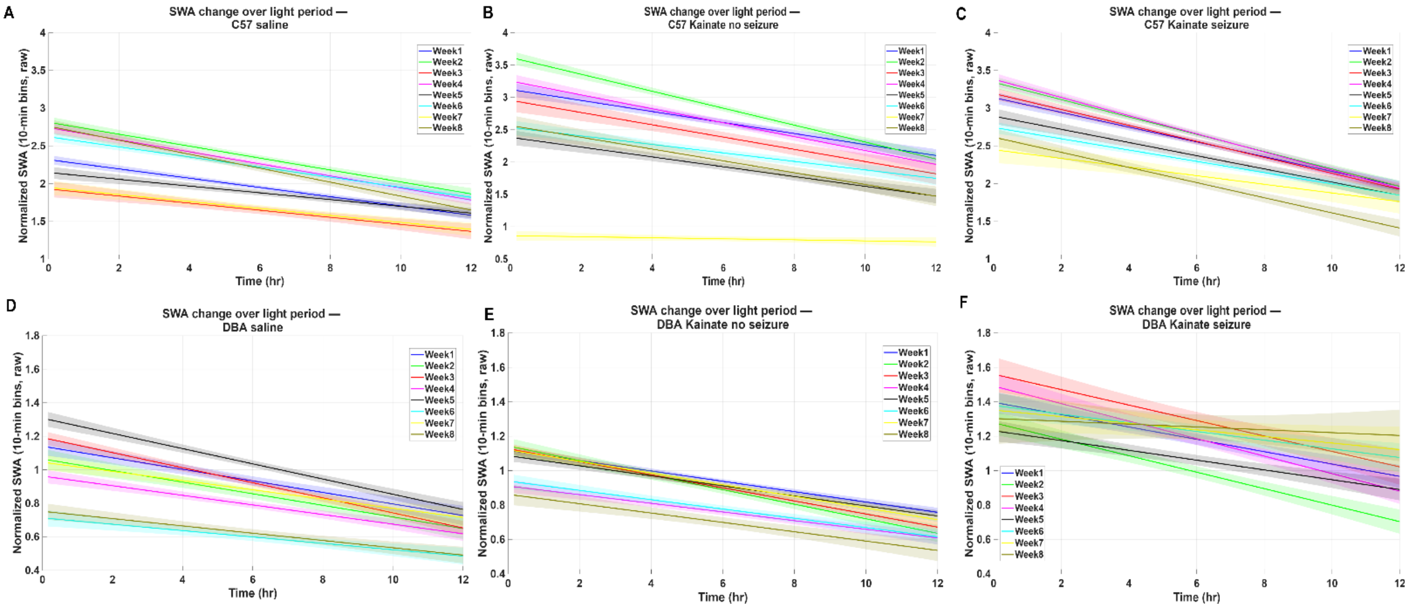


| **Table S1: Between group rhythm equality, MESOR, Amplitude and Acrophase differences between C57 and DBA before and after onset of seizures** | | |
|---|---|---|
| P Rhythm | C57 KA prior to seizure onset vs after seizure onset | **p<0.0001** |
| P Rhythm | DBA KA prior to seizure onset vs after seizure onset | p=0.303 |
| MESOR | C57 KA prior to seizure onset vs after seizure onset | **p<0.0001** |
| MESOR | DBA KA prior to seizure onset vs after seizure onset | p=0.105 |
| Amplitude | C57 KA prior to seizure onset vs after seizure onset | **<0.0001** |
| Amplitude | DBA KA prior to seizure onset vs after seizure onset | 0.517 |
| Acrophase | C57 KA prior to seizure onset vs after seizure onset | **0.047** |
| Acrophase | DBA KA prior to seizure onset vs after seizure onset | 0.195 |

| **Table S2: Change in slope of overnight decay of SWA from week-1 to week-8** | | | | | | | | |
|---|---|---|---|---|---|---|---|---|
| Treatment group | Week 1 | Week 2 | Week 3 | Week 4 | Week 5 | Week 6 | Week 7 | Week 8 |
| C57 Saline | -0.645 | -0.080 | -0.054 | -0.086 | -0.050 | -0.062 | -0.051 | -0.094 |
| C57 KA-seizures | -0.819 | -0.134 | -0.103 | -0.108 | -0.078 | -0.067 | -0.002 | -0.083 |
| C57 KA +seizures | -0.096 | -0.121 | -0.1066 | -0.123 | -0.093 | -0.083 | -0.064 | -0.108 |
| DBA Saline | -0.029 | -0.037 | -0.045 | -0.032 | -0.044 | -0.020 | -0.033 | -0.024 |
| DBA KA-seizures | -0.032 | -0.045 | -0.042 | -0.032 | -0.032 | -0.030 | -0.040 | -0.032 |
| DBA KA+seizures | -0.039 | -0.041 | -0.041 | -0.054 | -0.034 | -0.023 | -0.022 | 0.009 |
